# Lipid and nucleocapsid N-protein accumulation in COVID-19 patient lung and infected cells

**DOI:** 10.1101/2021.06.24.449252

**Authors:** Anita E. Grootemaat, Sanne van der Niet, Edwin R. Scholl, Eva Roos, Bernadette Schurink, Marianna Bugiani, Sara E. Miller, Per Larsen, Jeannette Pankras, Eric A. Reits, Nicole N. van der Wel

## Abstract

The pandemic of the severe acute respiratory syndrome coronavirus 2 (SARS-CoV-2) has caused a global outbreak and prompted an enormous research effort. Still, the subcellular localization of the corona virus in lungs of COVID-19 patients is not well understood. Here, the localization of the SARS-CoV-2 proteins is studied in postmortem lung material of COVID-19 patients and in SARS- CoV-2 infected Vero cells, processed identically. Correlative light and electron microscopy on semi- thick cryo-sections, demonstrated induction of electron-lucent, lipid filled compartments after SARS- CoV-2 infection in both lung and cell cultures. In lung tissue, the non-structural protein 4 and the stable nucleocapsid N-protein, were detected on these novel lipid filled compartments. The induction of such lipid filled compartments and the localization of the viral proteins in lung of patients with fatal COVID-19, may explain the extensive inflammatory response and provide a new hallmark for SARS- Cov-2 infection at the final, fatal stage of infection.

## Introduction

The outbreak of Severe Acute Respiratory Syndrome Coronavirus 2 (SARS-CoV-2) in late 2019 is the third major outbreak of β-coronaviruses in the human population of the past two decennia, together with the smaller outbreaks of Severe Acute Respiratory Syndrome Coronavirus (SARS-CoV-1) in 2003 and Middle East Respiratory Syndrome coronavirus (MERS-CoV) in 2012.

SARS-CoV-2 belongs to the family *Coronaviridae*, a large family of single-stranded positive-sense RNA ((+)RNA) viruses. The first two-thirds of the genome typically codes for polyproteins that, once processed by proteases, produce non-structural proteins involved in viral replication [1]. The remaining third of the genome consists of four structural proteins: envelope (E), membrane (M), nucleocapsid (N), and spike (S). Coronaviruses are well known for their ability to induce high membrane plasticity in host cells, where the membrane rearrangements lead to the formation of viral replication organelles (ROs) [2–10]. As observed in SARS-CoV-1, MERS-CoV, and the closely related coronavirus murine hepatitis virus (MHV), the ROs consist of convoluted membranes (CMs) that are interconnected with double-membrane vesicles (DMVs) and appear to be continuous with the membranes that constitute the endoplasmic reticulum (ER) [2,11–17]. Elaborate studies using immuno-fluorescence and electron microscopy (EM) techniques demonstrate that DMVs contain double-stranded RNA (dsRNA) which can be used as a marker of (+)RNA virus replication [2,3,18,19]. Taken together, these findings indicate that the RO serves as the replication and transcription site in which the DMVs, may provide a zone safe from detection by the innate immune sensors and degradation by RNA degradation machinery in the host cell [20, 21].

The formation of DMVs has been shown to be facilitated by coronaviral non-structural proteins (nsps) [22]. Co-expression of three virally encoded transmembrane proteins, namely nsp3, nsp4, and nsp6, has been found to be sufficient for the production of DMVs in SARS-CoV-1 and MERS-CoV where the interactions of nsp3 and nsp4 result in the pairing and curving of membranes, and nsp6 contributes to the production of vesicles [9,10,23]. A recent publication, using cryo-electron tomography (cryo- ET), shows DMVs of SARS-CoV-2 and MHV in a native host cellular environment containing pore complexes that were not found in previous studies using conventional EM methods [18]. Additionally, the publications by Wolff *et al.* 2020 [24] and Klein *et al*. 2020 [17] demonstrate the presence of N- protein in these DMVs.

The subcellular localization of the viral proteins and virus particles is based on infections in cultured cells. In patient material, viral proteins have been localized at a cellular level in various organs of COVID-19 patients [25], including human kidney [26], and in lungs of cynomolgus macaques [27]. These studies used light microscopy to find regions of interest, and some of the studies subsequently used EM to find virus particles. One of the hurdles to overcome is the correct identification of viral particles in patient material such as lung [12,28–32], kidney [33–39] and other organs reviewed in [6]. Recent publications show data on the morphology and size of isolated SARS-CoV-2 particles [40–43] and virus particles in Vero E6 cells [17] with the use of conventional EM and cryo-EM, although this data alone is not always sufficient to recognize viral proteins or virus particles. Bullock *et al*. proposed a set of eight rules for the correct identification of coronaviruses [6]. Following these rules, a closer inspection of 27 articles where supposed SARS-CoV-2 particles in patient-derived samples have been found, revealed that according to Bullock and Miller, only four articles correctly identified virus [6,44–47]. The most common misinterpretations were clathrin-coated vesicles as single SARS-CoV-2 particles and endosome-derived multi-vesicular bodies (MVBs) as ROs [6, 47].

To assist in this identification conundrum, labelling of antibodies directed against specific viral proteins can be of use. In this article, we provide the first insights into the localizations of both structural and non-structural proteins in SARS-CoV-2-infected Vero cells and compare this with identically processed patient samples retrieved during the first wave of SARS-CoV-2 infections using immuno-gold labelling and CLEM.

## Results

### Immuno-Electron Microscopy on SARS-CoV-2-infected Vero Cells

Since the outbreak of COVID-19, the identification of virus particles using EM in lung has been a heavily debated subject [6,48,49]. Based on the morphology, it is, especially in postmortem material, difficult to discriminate single virus particles from clathrin-coated vesicles, and MVBs have been interpreted as clusters of virus particles. Therefore, we decided to employ immuno-gold labelling, which can be used to decorate (viral) proteins specifically with 10- or 15-nm gold particles to distinguish them from cell organelles. This way, virus particles with M-, N-, or S-protein and the replication complexes with non-structural proteins can be identified by the gold attached to the specific antibodies. To validate whether the antibodies used for recognition of the proteins in FM [50] can be used on patient materials fixed with an extended fixation protocol, we first tested these antibodies on SARS-CoV-2-infected Vero cells. The antibodies were used on uninfected and 24-hour infected Vero cells fixed for 1, 3, and 14 days as we have fixed patient material in a similar manner. Different antibodies against viral proteins were tested (see Materials and Methods), and successful labelling and their subcellular localizations are described.

### Characterization of virus particles with N-protein

Immuno-gold labelling of SARS-CoV-1 structural proteins using a mouse anti-SARS-CoV-1-N (46-4) antibody demonstrated that the nucleocapsid protein (N-protein or N) is detected in the cytosol and on virus particles in several subcellular structures (Figs 1, S1) of infected cells. The N-protein can be specifically detected, as no labelling was detected on uninfected cells. Therefore, all N-protein positive, membrane enclosed spherical structures ranging in size from 60 to 120 nm in diameter and with an electron-dense core (e-dense, black) [6, 41], are annotated here as virus particles. Note that in cells and tissues stained with osmium and embedded in resin, membranes appear e-dense, whereas using the immuno-EM method on cryo-sections, membranes appear electron-lucent (e-lucent, white) [51]. This is due to the fact that with the immuno-EM method, membranes are not stained, but only surrounding proteins in the cytosol are stained with uranyl acetate. In 24-hour infected Vero cells, small clusters of N-protein can be detected in proximity to double membrane structures, bending around the N-protein cluster similar to that in the cryo-EM sections (Fig S1), [18]. Coronaviruses are known to be a membrane enveloped viruses, mostly detected inside host membrane structures [6, 52], and indeed the majority of the virus particles are surrounded by membranes [3, 17] (Figs 1A, 1B).

**Figure 1.**
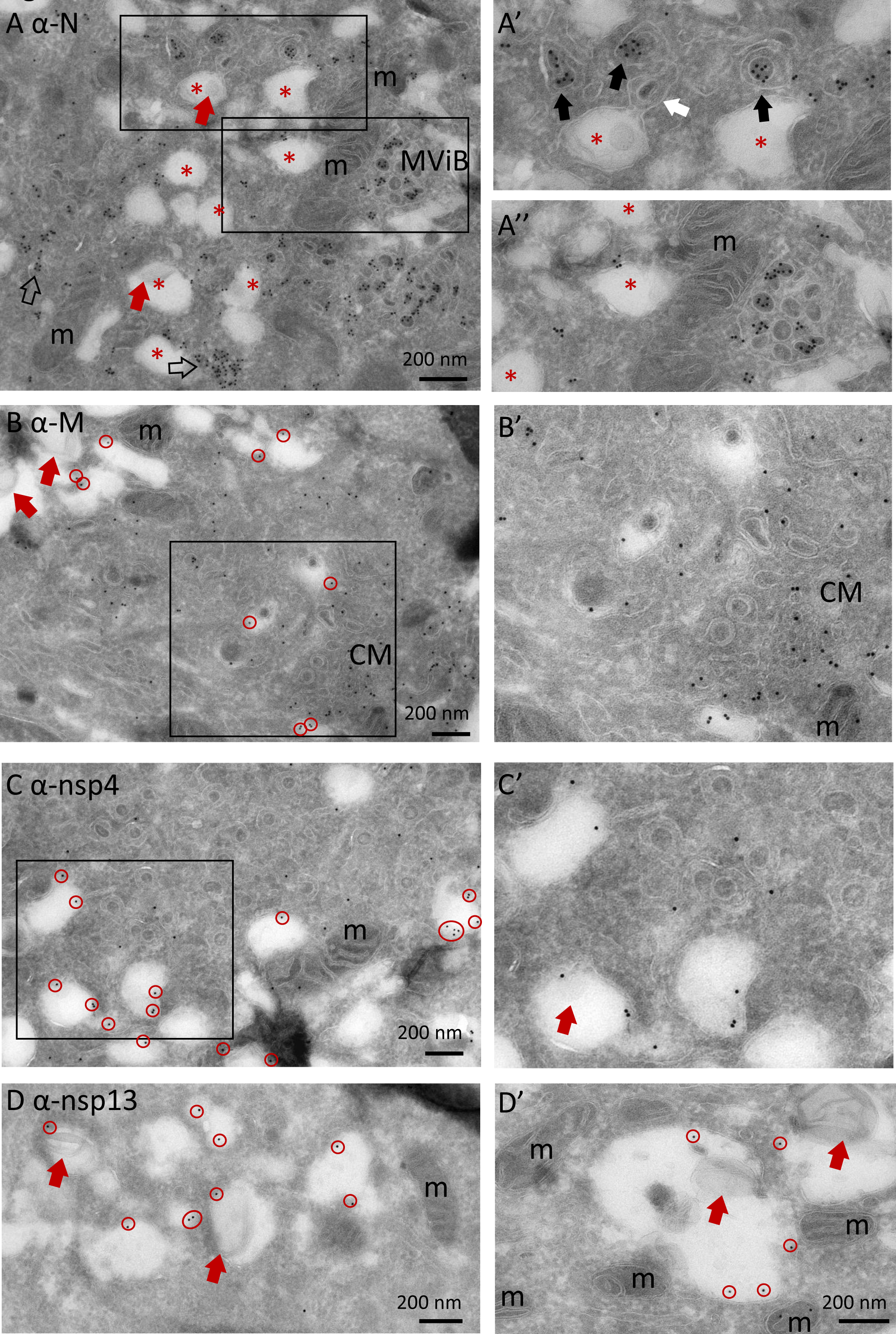
Subcellular localisation of viral proteins in infected Vero-cells. Vero cells were infected with SARS-CoV-2 for 24 hours and immuno-EM labelled with antibodies against SARS-CoV-1 proteins, followed by secondary antibodies conjugated to 10-nm gold particles. A) Clusters of N-protein labelling in cytosol (open arrows), and (enlarged in A’) on double membrane spherules (right- most black arrow), or virus particles enclosed in a single membrane (two left-most black arrows). From the e-lucent compartment (red *) a “virus-like” particle (as it is without N-protein labelling) is budding (white arrow). A’’) enlarged area with MViB containing labelled and unlabelled virus-like particles. B) M-protein immuno-gold labelling on e-lucent compartments (gold is circled in red); in enlarged box, immuno-gold labelling on convoluted membrane structure (CM). Note virus-like particles are not labelled. C) Immuno-gold labelling of nsp4 on e-lucent compartments (circled in red) and various virus like particles enclosed in a membrane without nsp4 labelling, also enlarged in C’. D) immuno-labelling of nsp13 on e-lucent compartments containing lipid like structures (red arrows). D’) higher magnification of D. Immuno-gold decoration on e-lucent compartments is indicated by red circles; mitochondria by m, multiple virus body by MViB, convoluted membrane structure by CM, lipid like structures by red arrows, N-protein in cytosol by open arrows, N-protein labelled virus by black arrows, and black boxes indicate enlarged area.

Intact viruses are also identified close to the Golgi (Fig S2), inside multi-virus bodies (MViB) (Fig 1A, S1, S3), inside open e-lucent structures (Fig S4). Both spherical and oval shaped virus particles are visible. The size of the virus particles are measured inside MViBs and intracytoplasmic and are categorized as spherical or oval-shaped. All particles are measured at the longest axis of N-protein-positive particles that have a clear membrane and e-dense core present. The average size between the spherical and oval-shaped virus is slightly different, but not statistically significant. Inside MViBs and intracytoplasmical, spherical particles are identical: 87 nm ± 17 nm versus 108 ± 27 nm and 113 ± 28 nm for the oval-shaped particles (Table 1). Different EM techniques result in slightly different sizes, being 97 ± 12 nm for oval-shaped cryo-EM fixed extracellular SARS-CoV-2 [43] or 99 nm in resin-embedded spherical virus [41]. Thus, in 24-hour infected Vero cells, N-protein-positive virus particles can be detected as spherical 87-nm to 113-nm oval-shaped membrane structures with an e-dense core, present in the cytosol, close to Golgi, or in multi-vesicular structures.

**Table 1.** Average particle size at different subcellular locations. Average size of virus particles in multi-virus bodies (MViB), and intracytoplasmic was measured and presented as average size (x) ± standard deviation and number of virus particles measured (n) in Vero cells infected with SARS-CoV- 2 for 24 hours and immuno-gold labelled for N-protein with 10-nm gold.

| Location | MViB |  | Intracytoplasmic |  |
| --- | --- | --- | --- | --- |
| Virus shape | spherical | oval | spherical | oval |
| x in nm | 87±17 | 108±27 | 87±17 | 113±28 |
| n | 61 | 21 | 59 | 8 |

### Classification of virus-containing compartments

As the presence of SARS-CoV-2 in multi-vesicular structures in lung is heavily debated [6, 48], we studied the presence of lysosomal markers like CD63 in the multi-virus bodies we showed to be N-protein positive (Fig 1, S1, S3). The Vero cell line is a kidney epithelial cell line from African green monkey, but antibodies against human CD63, a glycosylated transmembrane protein containing a putative lysosomal-targeting/internalisation motif, can be detected in multi-lamellar bodies (MLB) which are lysosomal compartments. Only some CD63 label is detected in the multi-vesicular bodies (Fig S3F). Therefore, we propose that the compartments in which the viral N-protein is detected, is not a true lysosome, but rather a multi-virus body. More elaborate studies on different stages of infection and blocking lysosomal acidification combined with immuno-EM have to be performed to determine the role of these MViBs during viral replication.

CD63 is also detected on early endosomes but not present on the majority of the e-lucent structures detected in clusters in SARS-CoV-2 infected cells (Fig S4). These structures seem to be induced by the virus infection, as uninfected cells contain larger lipid droplets but not the clustered e-lucent structures of 327 nm +/- 130 nm. High magnification analyses reveal that the e-lucent compartments appear to be filled with lipid like structures (Figs 1, S1B, S4F, S4H), much like we previously described for *Mycobacterium tuberculosis* infected cells [53]. Therefore, Nile red staining was performed on both uninfected and SARS-CoV-2 infected Vero cells, and a clear increase in Nile red signal is observed in infected cells (Fig 2). Indeed, others already demonstrated that lipid accumulation occurs after SARS-CoV-2 infection in Vero cells [54, 55]. To prove that the e-lucent compartments detected with EM are Nile red positive and thus lipid-containing compartments, both FM and EM were performed on the same section and combined in a CLEM image (Fig 2C). These CLEM images demonstrate that at least a part of the e-lucent compartments are lipid filled. The structure of these compartments is not identical to lipid droplets (LD), so we used an antibody specific for perilipin-2, which is known to localize in LD [56] to determine if the SARS-CoV-2 induced lipid filled compartments are in fact lipid droplets. Immuno-gold labelling is present on typical LD in uninfected Vero cells but not on the lipid filled compartments detected in SARS-CoV-2 infected cells (Figs 2F and 2G). Based on the absence of both the lysosomal marker CD63 and LD marker perilipin-2, these e-lucent structures are not lysosomes, nor LD but rather novel lipid-filled compartments induced by SARS-CoV-2 infection.

**Figure 2.**
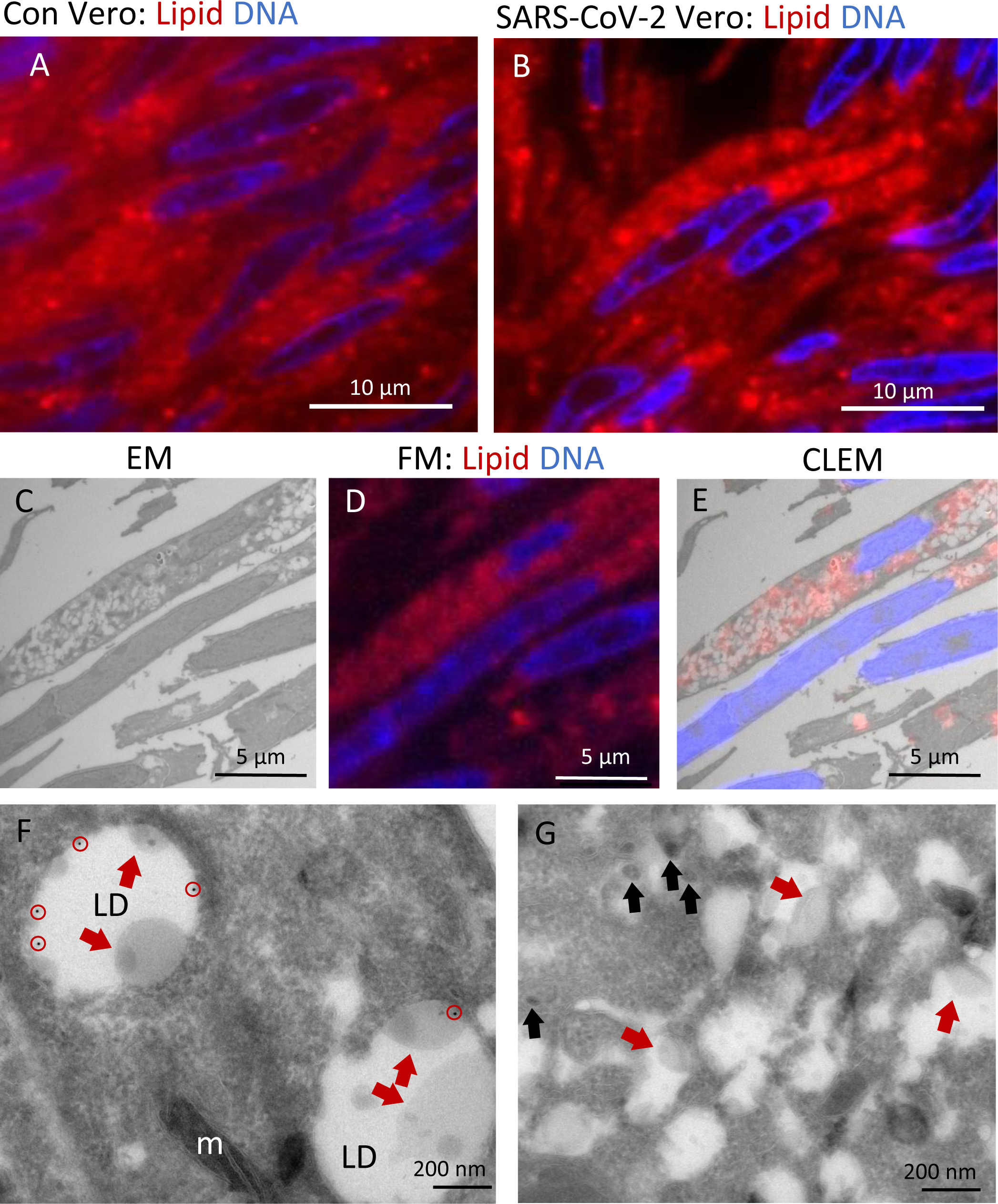
Lipid accumulates in e-lucent compartments more densely in infected Vero cells. Fluorescence microscopy of DNA and lipid staining with Nile red in A) the uninfected control (Con) Vero cells and B) cells infected with SARS-CoV-2 for 24 hours; C) Electron microscopy of infected cells; D) Fluorescence microscopy of the same cells, and E) Correlative light-electron microscopy (CLEM) showing lipid staining at e-lucent compartments in the electron microscope. Immuno-EM labelling for lipid droplet marker perilipin-2 in F) uninfected Vero cells and G) cells infected with SARS-CoV-2 for 24 hours. Blue color in A, B, D, and E shows the nuclei stained with Hoechst and red shows the lipids stained with Nile red. In electron micrographs, lipid like structure is denoted by red arrows, virus particles by black arrows, immuno-gold labelling of perilipin-2 by red circles, mitochondria by m, and lipid droplets by LD.

### Localisation of M-protein and non-structural proteins nsp4 and nsp13

The localisation of different viral proteins in cultured cells can be used to understand the pathology and replication of SARS-CoV-2 in lung tissue of COVID-19 patients. In infected Vero cells, the same procedures as for N-protein were applied to detect nsp3, but immuno-gold label is very limited, and thus, we conclude that this antibody does still recognize its substrate after 14 days of glutaraldehyde-paraformaldehyde fixation (Table 2). The non-structural proteins nsp4 and nsp13 are detected on vesicles located nearby and attached to the Golgi stacks (Fig S2). The signal of nsp13 is limited to a few gold particles per Golgi stack, and nsp4 is more distinct, but also has some background on mitochondria (Fig S4G). The M-protein abundantly labels Golgi stacks and vesicles around the Golgi. Interestingly, nsp4, nsp13, and M are also detected on MViBs (Fig S3) and at e-lucent lipid filled compartments, while uninfected cells are unlabelled (Figs 1, S4). These structures resemble double membrane vesicles (DMVs) or single membrane vesicles described for MHV, SARS-CoV-1, SARS-CoV-2, and MERS-CoV infected cells [3,5,16–18,23,50]. Single-membrane vesicles are proposed to be derived from the ER-to-Golgi intermediate compartment [57], and play a role in the secretion of virus to be released into extracellular space. With immuno-EM labelling only on some cellular compartments, a double membrane is detected (Fig S4H, blue arrows), which could be explained by the EM-technique used. Rather than performing high pressure fixation and freeze substitution [3] or cryo-EM [17, 18], we used conventional fixation to be able to compare Vero cells with lung tissues of COVID-19 patients. It is possible that the double membranes are lost during fixation for immuno-EM, as Snijder et al already demonstrated in 2006 [16]. Another limitation of the immuno-EM is that no clear spike proteins are detected on extracellular virus particles (Fig 3), though conventional sample preparation using osmium staining and embedding does show spikes [11, 41] as does cryo-EM [17, 18]. Extracellular virus particles are immuno-labelled for both N- and M-protein. Interestingly, the majority of the extracellular virus particles are not spherical, but rather oval-shaped. The subcellular localisation of N-, M-protein and nsps in infected Vero cells is summarized in Table 2, and translation of this knowledge to patient material could be essential for understanding COVID-19 pathogenesis in patients. As immuno-localisation with the antibodies against N-, M-protein, and nsp4, are specific and survive glutaraldehyde fixation, these antibodies can be used for analysis of lung tissues.

**Figure 3.**
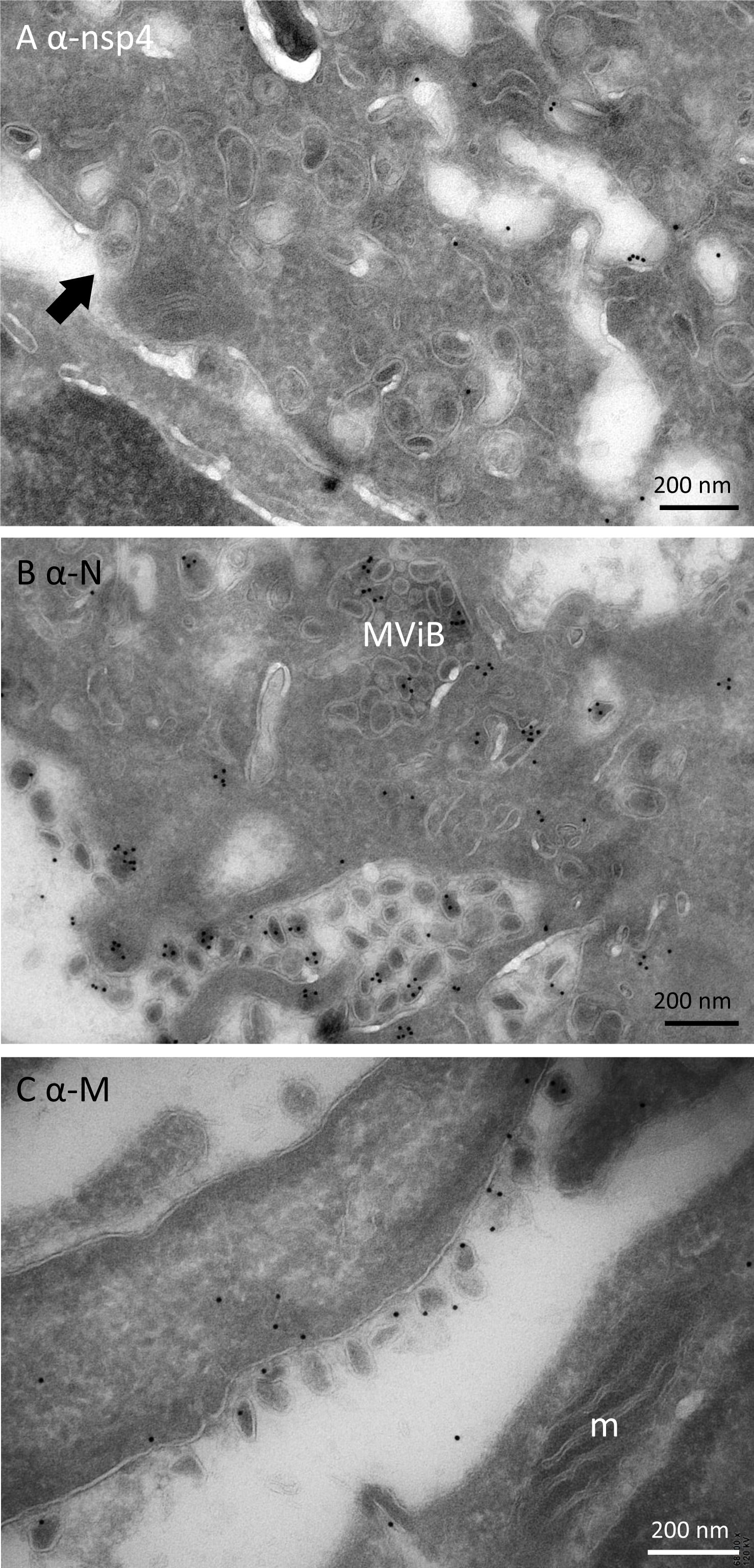
Release of virus particles from Vero cells infected with SARS-CoV-2 for 24 hr. EM micrographs, demonstrate A) lack of immuno-gold labelling on extracellular virus particle using anti- nsp4, a non-structural protein of SARS-CoV-2 (black arrow); B) extracellular virus particles labelled with anti-N-protein, and C) anti-M-protein also labels on extracellular virus particles. Here, m represents mitochondrion, MViB multi-virus body.

**Table 2.** Immuno-gold labelling of viral proteins in SARS-CoV-2-infected Vero cells. Presence of immuno-gold labelling on virus particles, Golgi, multi-virus bodies (MViB), multi-lamellar bodies (MLB), e-lucent compartments and extracellular virus particles in Vero cells infected with SARS- CoV-2 for 24 hours. Annotations: + present; - absent; +/- present but less prominent.

| Location in cell culture | N | M | Nsp3 | Nsp4 | Nsp13 | CD63 |
| --- | --- | --- | --- | --- | --- | --- |
| Virus particle | + | + | - | - | - | - |
| Golgi | +/- | + | - | + | +/- | - |
| MViB | + | + | - | +/- | +/- | +/- |
| MLB | - | - | - | - | - | + |
| e-lucent compartment | +/- | +/- | - | + | +/- | - |
| extracellular virus particle | + | + | - | - | - | - |

### Immuno-EM on lung of COVID-19 patients

In lung of COVID-19 patients, we searched for the presence of virus and replication organelles using antibodies selected on infected Vero cells. Material of 7 COVID-19 patients from a prospective autopsy cohort study performed at Amsterdam University Medical Centers (UMC) [25] were included.

With informed consent from relatives, full body autopsies were performed, and lung material was fixed for EM analysis. Materials were fixed for 1, 3 or 14 days. From those 7 patients, the lung tissues of 2 were too damaged to use for EM due to a postmortem delay. From our previous light microscopy analysis [25], we learned that only in a part of the lung tissue of a COVID-19 patient N-protein can be detected, and virus particles are difficult to find. Thus, in order to find the infected region of interest (ROI), we first performed fluorescence microscopy on sections of tissues processed for EM, so that when we identified a ROI containing viral proteins, EM could be performed (approached as in van Leeuwen et al., 2018). Semi-thick 0,3-µ m slices were incubated with antibodies against SARS-CoV-1 nsp3, nsp4, nsp13, and structural proteins N-, M-, and S-protein. We focused on areas near small blood vessels and alveolar walls, as our previous LM analysis revealed infected cells present along the alveolar walls. These cells were identified to be pneumocytes, stromal cells in the septa, endothelial cells in the septal capillaries, and alveolar macrophages [25]. Fluorescence microscopy showed that the N-protein (Fig 4) and nsp4 (Fig 5) could be detected. Noteworthy is the higher background for the M antibody and the relatively low labelling for nsp3 and nsp13 (Table 3).

**Figure 4.**
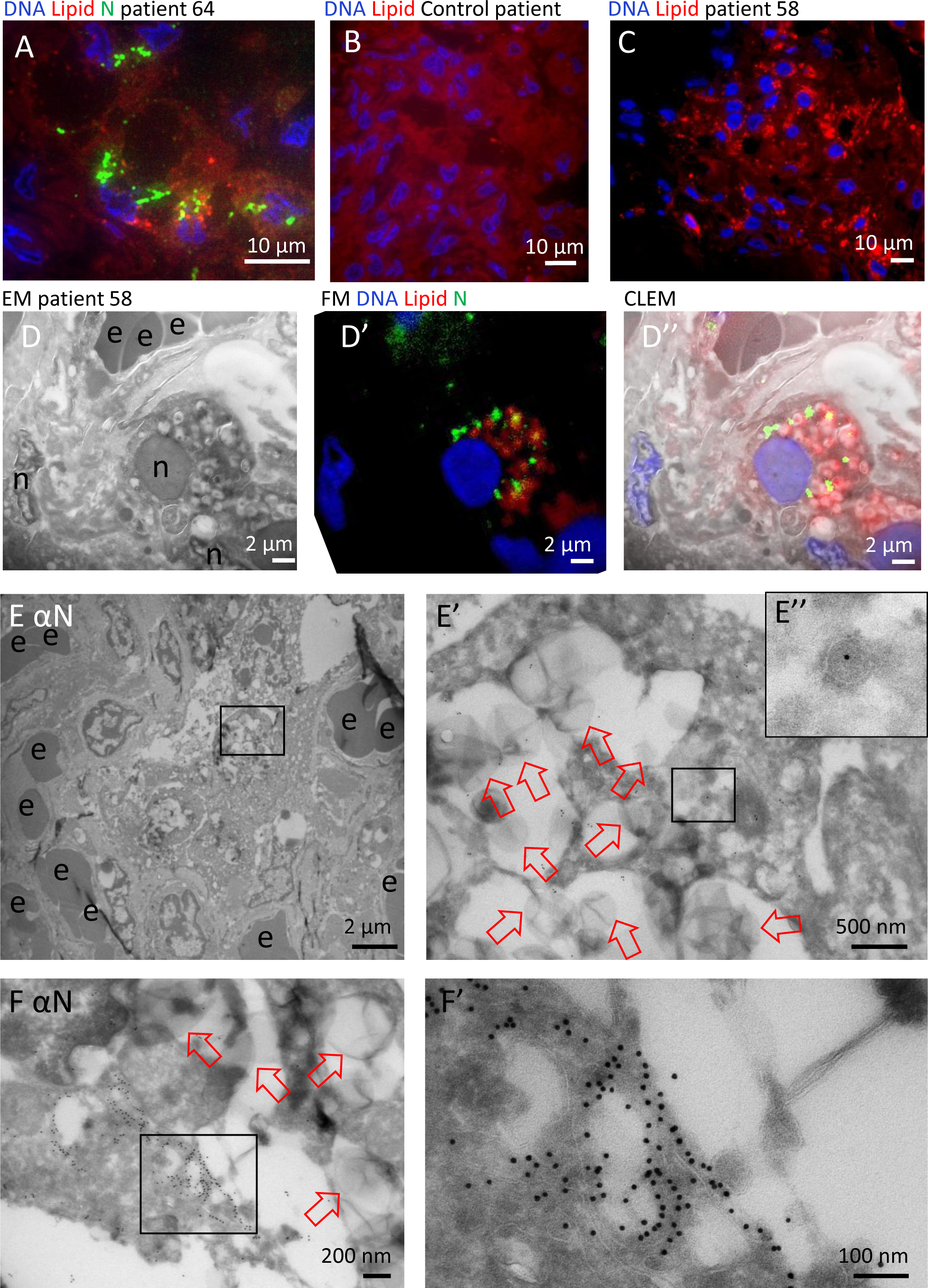
N-protein in e-lucent compartments in Lung COVID-19 patient. Lung from control and infected patients was either sectioned semi-thin for FM (A-C) or CLEM (D) and stained with Hoechst (blue) to identify nuclei, Nile red (red) to denote lipid, or anti-N-protein (green) to show N-protein, or it was ultrathin-sectioned for EM (E and F) and immuno-gold labelled using anti N-protein followed by secondary antibody tagged with 10-nm gold particles. A) COVID-19-infected lung showing accumulations of N-protein and Nile red stained lipids. B) overview of an uninfected control lung with no N-protein or lipid accumulation. C) Overview of infected lung with lipid accumulation. Identical section analysed by CLEM of infected lung demonstrate the e-lucent compartments present by EM (D) are Nile red and N-protein labelled (D’) by the overlay of the FM on the EM micrograph (D’’). Immuno-gold labelling of infected lung with antibody against N-protein at low magnification (E) and magnified region from boxed area where lipid like structures (open red arrows) are visible (E’) and a single virus particle with N labelling (E’’), low magnification of N-protein labelling on membrane structures near the e-lucent compartments F); high magnification of F’) clusters of N-protein labelling. Erythrocytes represented by e, nucleus by n, open red arrow lipid like structures, and boxed areas enlarged region.

**Figure 5.**
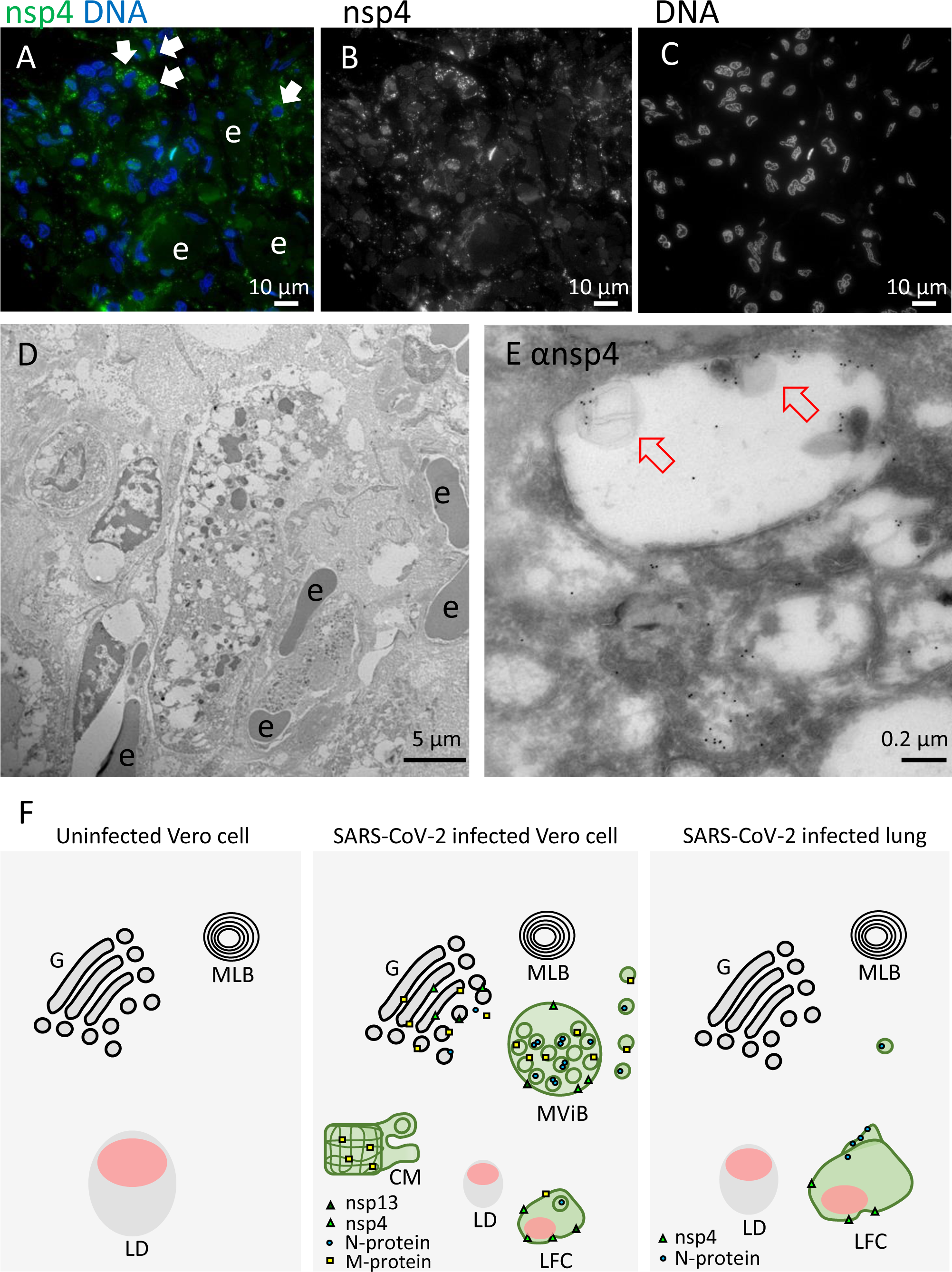
Non-structural protein 4 in e-lucent compartments infected lung. Lung tissue of COVID-19 patient 58 was either sectioned semi-thin for FM with A) nuclei, stained with Hoechst (blue), nsp4 stained with Alexa (green) in nsp4 positive cells indicated by white arrows and in black and white, and erythrocytes represented by e. Separate channels of nsp4 (B) and DNA (C). Ultrathin sections of infected lung immuno-gold labelled against nsp-4 and 10-nm gold particles in overview (D) and at higher magnification (E) e-lucent compartments with nsp4 labelling on membrane and lipid like structures (open red arrows) erythrocytes represented by e. F) Schematic representation of uninfected Vero cells, SARS-CoV-2 infected Vero cells and lung tissue of COVID-19 patient summarizing presence cellular organelles and subcellular localisation viral proteins. In black: host compartments, in green: viral compartments, in pink: lipid like structures, CM convoluted membrane G: Golgi, LD: lipid droplet, MLB: multi-lamellar bodies, MViB: multi-virus body, LFC: lipid-filled compartment and immuno-labelling viral proteins: dark green triangle: nsp13, light green triangle: nsp4, blue circle: N-protein, yellow square: M-protein.

**Table 3.** Immuno-gold labelling of viral proteins in SARS-CoV-2 infected lung. Presence of immuno-gold labelling on virus particles, Golgi, multi-virus bodies (MViB), multi-lamellar bodies (MLB), e-lucent compartments and extracellular virus particles in patient 58 and 64 infected with SARS-CoV-2. Annotations: + present; - absent; +/- present but less prominent.

| Location in lung | N | M | Nsp3 | Nsp4 | Nsp13 | CD63 |
| --- | --- | --- | --- | --- | --- | --- |
| Virus particle | +/- | - | - | - | - | - |
| Golgi | - | - | - | - | - | - |
| MViB | - | - | - | - | - | - |
| MLB | - | - | - | - | - | + |
| e-lucent compartment | +/- | +/- | - | + | - | - |
| Extracellular virus particle | - | - | - | - | - | - |

Thus, in lung tissue from COVID-19 patients, an ROI was selected by FM using the N-protein antibody. In one patient (patient 64), relatively large clusters of N-protein were detected (Fig 4A) often in a perinuclear region. As in Vero cells (Fig 2), an increase in lipid accumulation, was observed (Figs 4B, 4C). Nile red staining was combined with N-protein labelling, and N-protein and lipid accumulations, localize in the same general areas but did not co-localize at the same subcellular localisation. Control lung material processed identically to COVID-19 patient material and tested for lipid accumulation demonstrated homogeneous background staining. Sections of 150 nm were analysed with both FM and EM and combined (Fig 4D). In line with our CLEM data on the Vero cells, performing CLEM on lung tissue demonstrated that lung tissue also accumulates lipid in e-lucent compartments. Then ultrathin 60-nm cryo-sections were cut, and protein A conjugated to 10-nm gold particles was used to label N-protein.

The ultrastructure of the lung tissue is reasonable, given the fact that this is postmortem material and that it is from a patient with COVID-19. The tissue is unlike healthy lung tissue, not ventilated, but instead filled with erythrocytes and packed with inflammatory cells infiltrating the alveolar lumen and inter-alveolar septa. It is not always possible to identify the cell type specially when the nucleus is not present in the 60-nm thin section. N-protein is detected in cells with large e-lucent compartments, with some label found in e-lucent, lipid filled compartments. Only a few spherical single membrane structures with N-protein were detected (Fig 4 inset). These might be virus-like particles, but due to the low labelling (1 gold particle), the on average larger diameter (110 nm), and an atypical localisation in the cytosol, over-interpretation is possible. Nonetheless, large clusters of viruses are not detected. Besides the limited labelling on small round vesicles, N-protein is also present on membranous structures close to e-lucent compartments (Figs 4F, S5). These structures are not present in all patients; from the 7 patients investigated, 2 had clusters of proteins detectable with the SARS-CoV-1 anti N-protein. In patient 64 (patient description in Schurink et al., 2020), relatively large N-protein clusters at the e-lucent compartments were detected (Figs 4F, S5A-D), and smaller clusters are detected in patient 58, albeit at a similar location [on membrane clusters near the e-lucent compartments (Figs S5E, S5F)].

Using FM, nsp4 was identified in the same ROI of lung tissue used for detection of N-protein (Fig 5). Cells positive for nsp4 are present in various tissue compartments. Although background labelling is detected, some cells are brightly positive. Immuno-EM demonstrates nsp4 on e-lucent compartments, which are filled with lipid like structures. A small amount of label is detected on mitochondria which should be regarded as background labelling, as this is also present on uninfected Vero cells (Fig S4G). The summary of subcellular viral protein localisation in lung is presented in Table 3 and, compared to the quantity of labelling in Vero cells, less labelling is detected in only limited compartments. The lipid filled compartments, however, are positive for nsp4, and N-protein is accumulated close to these compartments. Like in Vero cells, lysosomal marker CD63 is absent from these compartments and thus the lipid filled compartments in lung are non-lysosomal. To our knowledge, these lipid filled compartments, containing viral proteins nsp4 and N-protein, have not been identified before and need to be further characterized.

## Discussion

Since the outbreak commenced, the identification of corona viruses in lung by EM has been debated, and several articles had to be revised [47,59,60]. Experienced Electron Microscopists [6] have summarized these studies and suggest using one of 3 strategies: 1) visualisation of viral morphogenesis, 2) immuno-EM or *in situ* hybridization, or 3) visualization of particles *in situ* in tissue combined with biochemical evidence of viral presence. We chose immuno-EM with gold labelling using already validated antibodies raised against SARS-CoV-1 [50]. Immuno-EM on Vero cells identified the monoclonal anti-SARS-CoV-1-N 46-4 to be the best for the detection of nucleocapsid N-protein. Virus particles were detected in the process of development as denoted by clusters of cytosolic N-protein surrounded by double membranes (Figs 1, S1). Spherical and/or oval virus particles are detected in MViBs, and in membrane clusters in the cell. The spherical virus particles are stable in size (87 ± 17 nm) and the oval-shaped virus particles are slightly larger (108 ± 27 in MViBs and 113 ± 28 nm for intracytoplasmic) than the spherical ones albeit, these variances are not statistically different. It should be noted that in immuno-EM and at 24 hours of infection, 20 % of the virus particles are scored as oval. The functional difference between spherical versus oval-shaped virus particles still has to be discovered but others have demonstrated that the oval or ellipsoidal- shaped virus particles contain more complexes of RNA and N-protein [43].

In lung of patients who had a fatal COVID-19 infection, virus-like particles are rarely detected even though the N-protein is detected in close proximity of the viral induced lipid filled compartments. In Vero cells however, N-protein is detected inside virus particles. It is possible that the difference is caused by incomplete fixation of lung or that ultrastructure is deteriorated in postmortem material. The overall ultrastructure of the tissue, however, is acceptable (Figs 4, 5), because the postmortem time was kept to a minimum and lung tissue was fixed within a few hours, during the first wave of COVID- 19 infections in the Netherlands. Finally, it is important to note that the magnification of EM makes finding 90-nm sized virus particles in a tissue block of 1x1 mm^2^ , extremely difficult. Still, some studies have detected an occasional cell filled with virus-like particles [44–47].

Interestingly, our CLEM data (Figs 2 and 4) demonstrated that part of the e-lucent compartments we have detected in Vero cells and in lung of COVID-19 patients are lipid-filled. Lipids are notoriously difficult to fix with glutaraldehyde and paraformaldehyde alone [61], and thus part of the compartments might have lost the lipid content but lipid accumulation in virus induced compartments is extremely interesting. For viruses of the Flaviviridae family, such as the dengue virus, hepatitis C virus and others, lipid accumulation has been shown to be involved in viral replication [62–69]. High resolution EM studies on cryo-preserved MHV infected cells, suggest DMVs to be filled with viral RNA with LD lying next to the DMVs [18].Also in infected human pulmonary epithelial Calu-3 cells [13] lipid droplets are detected close to the DMVs. Fluorescence microscopy studies have demonstrated lipid accumulations in SARS-CoV-2-infection, in Vero cells [54] and Nardacci et al., 2021, demonstrated that lipid accumulation is specific for SARS-CoV-2 and not for SARS-CoV-1 in a comparative electron microscopy study and established an increase of LD in lungs from deceased COVID-19 patients. Here, using immuno-EM, combined with Fluorescence microscopy, demonstrate an induction of lipid filled compartments and propose the SARS-CoV-2 infection-induced compartments are not LDs, as they are irregular in shape and have a different morphology than spherical perilipin-2 stained LDs. Also the clearly visible membrane (Fig S2), containing transmembrane proteins nsp4 and nsp13, demonstrates that the viral induced lipid-filled compartments are surrounded by a bilayer, while lipid droplets are surrounded by a monolayer of phospholipids.

Taken together, SARS-CoV-2 infection induces novel lipid filled compartments, different from LD or endosomes but with viral proteins nsp4 and N-protein.

Here we used the SARS-CoV-2 infected Vero cells to determine in which fixation conditions and on which structures we could detect viral proteins, to compare that with lung tissues from patients conserved in the same conditions. We noticed that not all structures present in Vero cells can be detected at the last stage of infection. A virus induced structure that is well described is convoluted membranes, which was detected in Vero cells (Fig. 1B) but not in lung at the final state of infection. In addition, multi-virus bodies were specifically detected in Vero cells and not in lung (Figs 1, S1, S3).

The MViBs are different from lysosomal MVBs, based on the fact that the MViBs are not CD63 positive and based on the size, morphology and the M- , N-protein labelling detected within the structures. In the lung of patients with fatal COVID-19, no MViBs were detected. Also double membrane vesicles (DMVs), have been described in several EM studies [2,3,5,15–18] but are not so obvious in our immuno-EM images; only a few double membranes were identified surrounding e- lucent compartments (Fig S4, blue arrows), possible due to fixation limitations, as shown before by Snijder *et al.,* 2006 [16]. As double membranes were not recognizable, DMVs were not annotated in this study. Recent comparison of SARS-CoV-2 infected Vero cells versus lung organoids demonstrated that the subcellular trafficking in Vero cells might be different [70] which can explain the presence of MViB in Vero and absence of these organelle in lung. Also the infections stage could be an explanation as we have analysed postmortem material and thus the last stage of the disease.

Remarkably, N-protein and nsp4 are detected in lung of patients in the last stage of the disease. It seems unlikely that only these 2 proteins are still produced by active replication of the virus, rather both N-protein and nsp4 are more stable proteins and thus not degraded. The gene encoding the N-protein is conserved and stable, and the N-protein itself is both highly immunogenic and highly expressed during infection [71]. Work on patients with a SARS-CoV-1 infection demonstrated elevated levels of IgG antibodies against N-protein [72] and showed that N-protein is an antigen for T-cell responses, inducing SARS-CoV-1-specific T-cell proliferation and cytotoxic activity [73–75].

Also, in an increasing number of case studies, anti-N IgGs were detected in patients with severe COVID-19 [76] and in children, 5 out of 6 produced neutralizing IgG and IgM antibodies targeted to the N- and S-proteins of SARS-CoV-2 [77]. Interestingly, recent reports show that immune responses to the N-protein have been associated to poor clinical out-comes [78] and correlates with severity of COVID-19 [79].

In the current electron microscopy study, we detected in fatal COVID-19 infections using SARS-CoV-specific antibodies, the stable presence of N-protein and nsp4 on novel lipid filled compartments.

Already, it has been demonstrated pharmacological inhibition via a key enzyme for LD formation effected SARS-CoV2 replication cells [52] suggesting that lipid accumulation is a potential drug target. The identification of the lipid filled compartments could serve as a hallmark for SARS-CoV-2 infections, especially since finding virus particles is challenging. Also we speculate that lipid-filled viral protein-containing compartments play an important role in the secondary effects of the disease. The uncontrolled immune responses causing the devastating damage of COVID-19 likely is responding to either the proteins or even lipids accumulating in these novel subcellular compartments and thus reducing lipid accumulation, will provide new therapeutic strategies.

## Materials and Methods

### EM Infection and fixation of Cultured Vero Cells

Vero E6 were seeded (2.5x10^6^ cells/T75 flask) one day before infection in MEM/25mM HEPES/2% fetal calf serum with penicillin and streptomycin. Cells (∼5x10^6^ cells/T75) were infected with MOI=0.2 by adding the virus (nCoV-2019/Melb-1, (4.3x10^6^ pfu/ml) to each T75 flask. Incubation was performed at 37°C for 24 hours. Then cells with and, as a control without virus, were fixed in 1 part medium plus 1 part 6% PFA + 0,4% GA in 0,4M PHEM buffer (240mM Pipes, 100mM HEPES, 8mM MgCl_2_ and 40mM EGTA at pH 6.9). After 1, 3 and 14 days of fixation samples were transferred to storage buffer (0,2M PHEM with 0,5% PFA).

### Collection and initial fixation of tissue from COVID-19 patients

Autopsies were performed at Amsterdam University Medical Centers (UMC), at the VU Medical Center, and the Academic Medical Center, the Netherlands, according to the declaration of Helsinki. For this EM study, 7 patients with clinically confirmed COVID-19 for whom autopsy was requested, were included (Table 4). Ethical approval was granted by the institutional review board of Amsterdam UMC (METC 2020.167). As described by Schurink et al., 2020 COVID-19 was confirmed by quantitative real-time RT-PCR, and informed consent was obtained from the decedents’ next of kin.

**Table 4.** Patient description. Information of patients from who autopsy material was taken with informed consent and fixed for electron microscopy. In this study, electron micrographs were used from patients SVU 20-58, SVU 20-64, and control T18-5683.

| Patient | Infection stadium | Sex | Age | COV-N | Remarks |
| --- | --- | --- | --- | --- | --- |
| SVU 20-58 | Limited infected cells in lung, limited systemic presence (HPB tract) | F | 72 | + | Data presented |
| SVU 20-39 | Severe infected cells in lungs, systemic presence (GI tract) | M | 73 | + |  |
| SVU 20-63 | No presence in lung, limited presence in the heart | M | 74 | + |  |
| SVU 20-64 | Limited presence in the lung, no systemic presence | F | 68 | + | Data presented |
| SVU 20-155 | - | F | 75 | + |  |
| SVU 20-163 | - | M | 61 | + |  |
| SVU 20-174 | - | M | 78 | + |  |
| SVU 20-129 | Control non-covid | M | 68 | - |  |
| T18-5683 | Control non-covid | F | 5 | - | Data presented |
| T18-10645 | Control non-covid | F | 15 | - |  |

During autopsy, lungs for conventional EM were fixed in Karnovski fixative with 4% PFA with 1% GA in 0,1 M sodium cacodylate buffer. To avoid safety problems, samples were fixed for 14 days and transferred to storage buffer or embedded in gelatin and snap frozen.

### Embedding and sectioning

After fixation, cells and tissue were washed 3 times with phosphate buffered saline (PBS) + 0.02M glycine (Merck, K27662101) to remove fixative. Cells were pelleted by centrifugation at 980 xg for 3 minutes. Supernatant was removed, and cells were directly embedded in 12% gelatin (Sigma, G2500- 500G) in 0.1 M phosphate buffer and pelleted by centrifugation for 3 minutes at 10,950 xg and solidified on ice, and blocks of ∼1 mm^2^ were cut with a razor blade. Lung tissue was cut into blocks of 1-2 mm^2^ and imbedded in a gelatin series of 2%, 6%, and 12% gelatin in 0.1 M phosphate buffer.

Blocks of cells or tissue were incubated overnight in 2.3M sucrose at 4°C (Merck, K17687153) in 0.1M phosphate buffer. Then samples were snap frozen and stored in liquid nitrogen. Sectioning was performed using a diamond knife (Diatome cryo-immuno) on a Leica Ultracut UC6 cryo-ultramicrotome. Semi thin sections (150-300 nm) were made at -80°C, and ultrathin sections were made at -120°C. The sections were transferred to a formvar-coated copper grid, gold finder grid, or glass slide in a droplet of 1 part 2% methylcellulose (Sigma, M6385-250G) to 1 part 2.3M sucrose. Sections were stored at 4°C until labelling.

### Immuno-fluorescence labelling

Semi-thin cryo-sections were transferred to gold finder grids for EM or to glass slides for light microscopy (LM) and washed with PBS + 0.02M glycine. Then, for LM, semi-thin sections were incubated on primary antibody for 1 hour in PBS + 0.1% bovine serum albumin (Sigma, A4503-50G) and washed with PBS + 0.02M glycine. Thereafter, they were incubated with secondary antibody conjugated to Alexa 488 (Mol. Probes, A32731), and in the last 5 minutes, Nile red (Sigma, 72485) and Hoechst 33342 (Thermo Fisher, H3570) was added. After washing with PBS, a cover slip was mounted with Vectashield (Vector laboratories, H-1000). Glass slides were imaged using a Leica DM6 widefield microscope with a 100x oil objective. Images were analyzed using ImageJ FIJI.

### Immuno-gold labelling

For EM, ultrathin sections were picked up and placed on 150 mesh copper grids and incubated on 2% gelatin in 0.1M phosphate buffer for 30 minutes at 37°C. Then, at room temperature, grids were washed with PBS + 0.02M glycine and blocked with 1% BSA in PBS. Grids were incubated with primary antibody in 1% BSA in PBS for 45 minutes. Then, grids were washed with PBS + 0.02M glycine. When the primary antibody was an unlabeled mouse monoclonal antibody, a secondary antibody, raised against mouse serum was used as a bridge to enhance labelling, followed by incubation with protein A conjugated with colloidal gold. In this case, background blocking was done by 0.1% BSA in PBS + 0.02M glycine, followed by incubation on rabbit anti mouse antibody (Z0259, DAKO) for 20 minutes and washed with PBS + 0.02M glycine. Again, grids were incubated in blocking solution and subsequently with protein A conjugated to 10-nm gold (Utrecht University).

After washing with PBS, grids were incubated with 1% glutaraldehyde in PBS to fix the antibody-gold complex and washed 10 times for 2 minutes each with water. To contrast the samples, grids were incubated with uranyl acetate in 2% methylcellulose for 5 minutes, and the excess liquid was blotted from the grids with filter paper. Grids were imaged using a FEI Tecnai 120kV transmission electron microscope with a Veleta or Xarosa camera (EMSIS). Images were analyzed using imageJ FIJI.

### Correlative light and electron microscopy

For CLEM, we used a method described earlier [58]. In short; grids were washed with PBS + 0.02M glycine and incubated for 1 hour with primary antibody and again washed with PBS + 0.02M glycine. Thereafter, grids were incubated with secondary antibody Alexa 488 and in the last 5 minutes Nile red (Sigma, 72485) and Hoechst 33342 (Thermo Fisher, H3570) were added. After washing in PBS, the grids were mounted in between a glass slide and a coverslip in a droplet of Vectashield. CLEM samples were imaged on a Leica DM6 widefield microscope using a 100x oil objective. Images were analyzed using LasX. After widefield imaging, the coverslip was removed from the glass slide by pipetting PBS in between the coverslip and the glass slide. Vectashield was removed by washing the grid with milliQ water at 37°C. Thereafter, the grids were contrasted and imaged as described above. The correlation was performed using ICY eC-CLEM software.

## List of materials

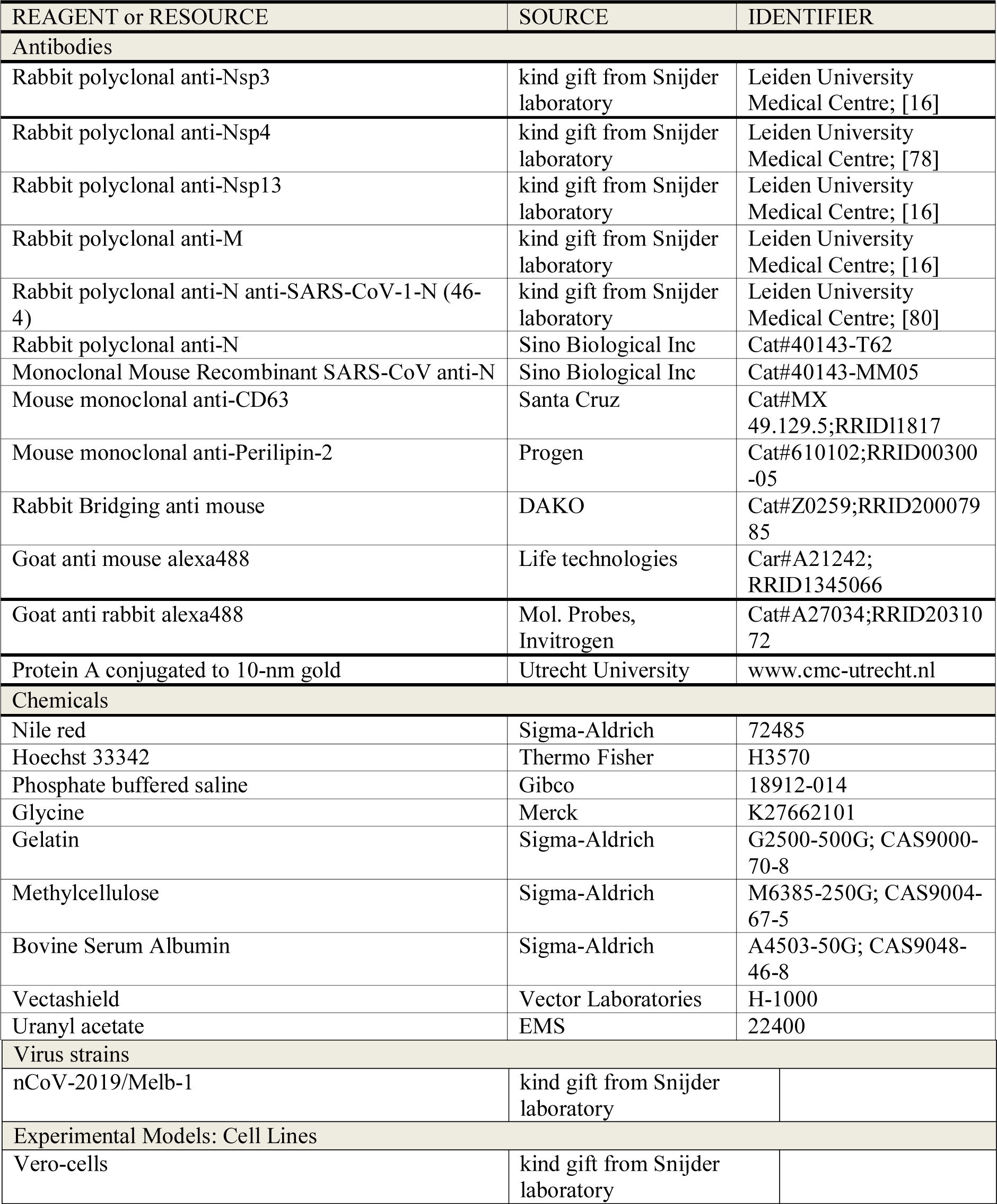

## Supporting information

Supplemental Figures Grootemaat

## Acknowledgements

We like to thank Eric Snijder, Montse Barcena for input, discussion and providing SARS-CoV-2 infected Vero cells, Sabine Krom and Jordy de Bakker for technical assistance, Sandrine Florquin for providing control lung materials; ER, MB thank Funding Amsterdam UMC Corona Research Fund; SvdN was funded by NADP and NIH grant no. AI116604.

