## Supplemental Figures Grootemaat for "Lipid and nucleocapsid N-protein accumulation in COVID-19 patient lung and infected cells"

Supplemental Figure 1

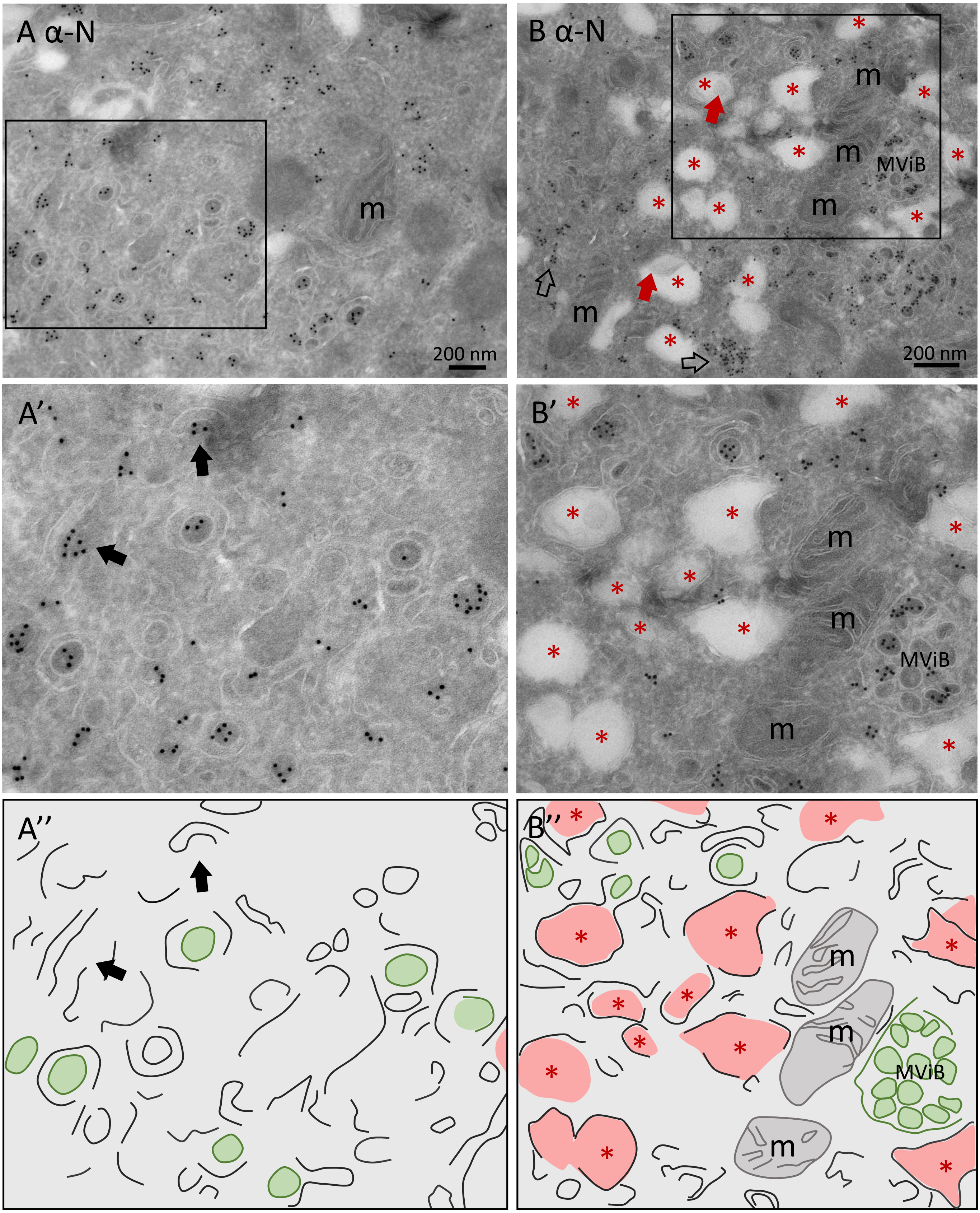

**Supplemental Figure 1. Intermediated stages of SARS-CoV-2 virus formation.**

Electron micrographs with detail of cytosol of 24-hour-infected Vero cell with immuno-gold labelling of N-protein and 10-nm gold particles with A) overview part cytosol with virus particles and in A') boxed area in higher magnification with double membranes surrounding a cluster of cytosolic N-protein indicated by black arrows and in A")a colour coded overlay for interpretation. B) same electron micrograph as in Figure 1, with clusters of N labelling in the cytosol indicated by open arrows, lipids like structures inside e-lucent compartments by red arrows, and boxed area in higher magnification in B') and a colour coded overlay for interpretation in B"). Colour coded overlay of A" and B" indicate in green virus particles with e-dense core and anti-N-protein immuno-gold labelling, in black lines host membranes, e-lucent compartments in pink with red *, mitochondria in grey with m and multi virus body by MViB.

Supplemental Figure 2

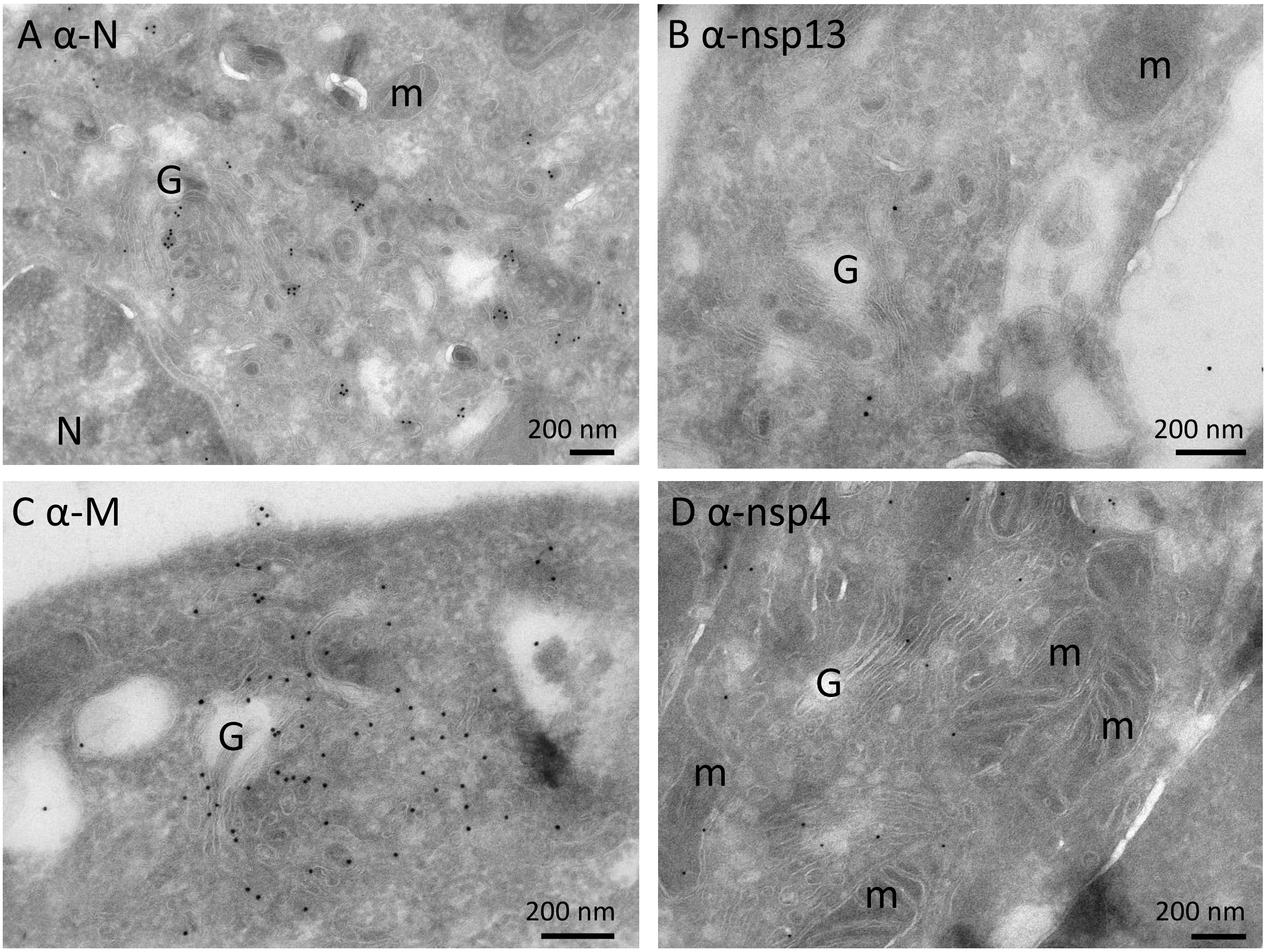


**Supplemental Figure 2. Immuno-gold labelling near a Golgi stack.** Electron microscopy of Vero cells infected with SARS-CoV-2 for 24 hours and immuno-labelled with antisera and 10-nm gold particles. A) N-protein labelling on virus-like particles near a Golgi stack. B) A few nsp13-labelled vesicles close to a Golgi stack. C) Abundant M-protein labelling on the membrane of a Golgi stack and virus-like particles near the Golgi stack. D) nsp4 labelling on membranes of a Golgi stack and on mitochondria. Mitochondria represented by m, Golgi by G, nucleus by n.

Supplemental Figure 3

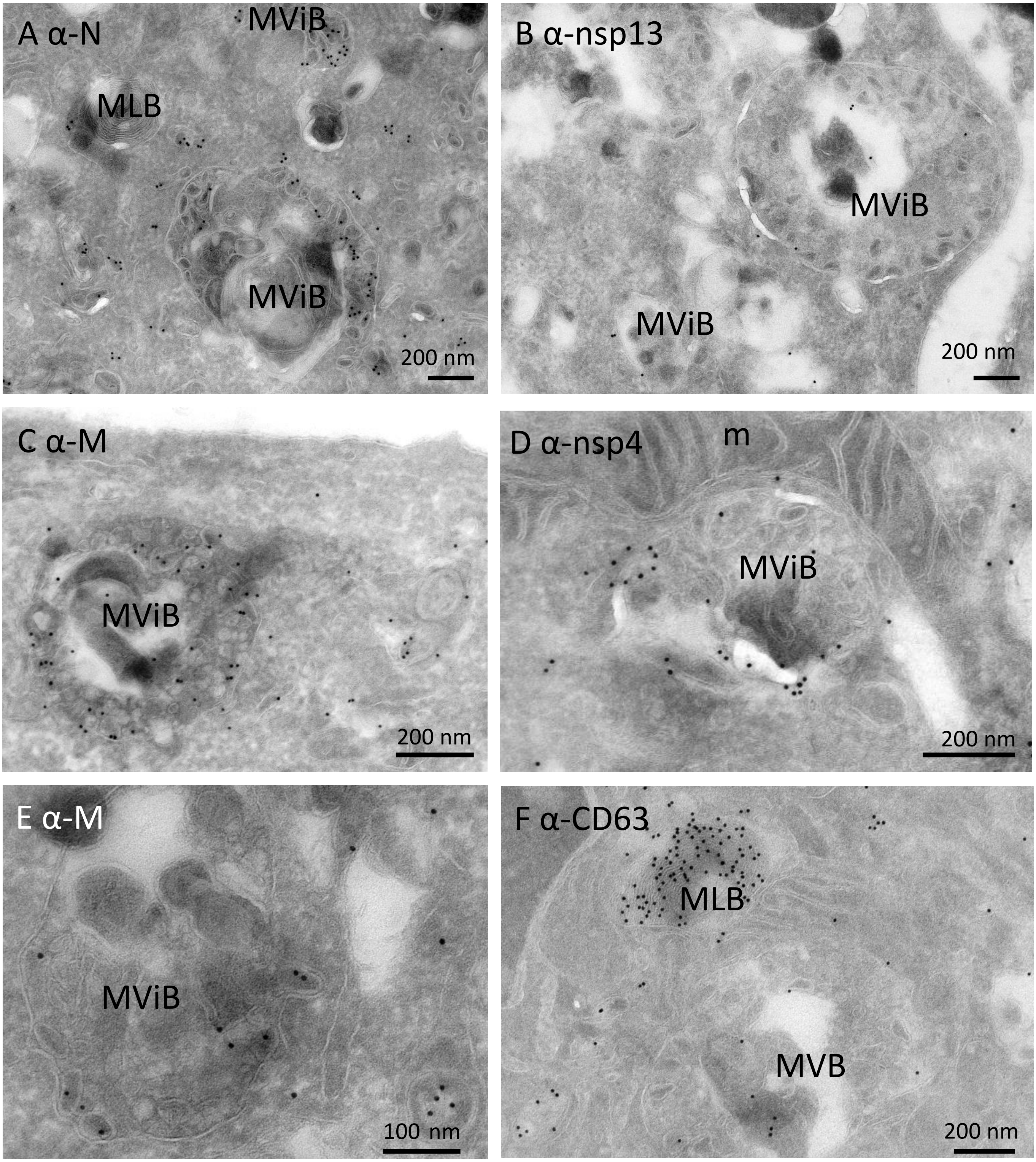

**Supplemental Figure 3. Presence of viral proteins in multi-virus bodies.** Electron micrographs of Vero cells infected with SARS-CoV-2 for 24 hours and immuno-gold labeled with antisera raised against SARS-CoV-1. A) N-protein labelling on virus-like particles in an MViB compartment, and not in an MLB, B) some nsp13 labelling on the MViB. C) Abundant M-protein labelling on virus-like particles in the MViB. D) nsp4 labelling on an MViB, and E) M-protein labelling on an MViB with a virus particle pushing the limiting membrane, F) CD63 labelling as a lysosomal marker is detected on the MLB but less on the multi-vesicular body (not MViB, as no viral proteins detected). Multi-virus body represented by MViB, multi-vesicular body by MVB, multi-lamellar body by MLB, mitochondrion by m.

### Supplemental Figure 4

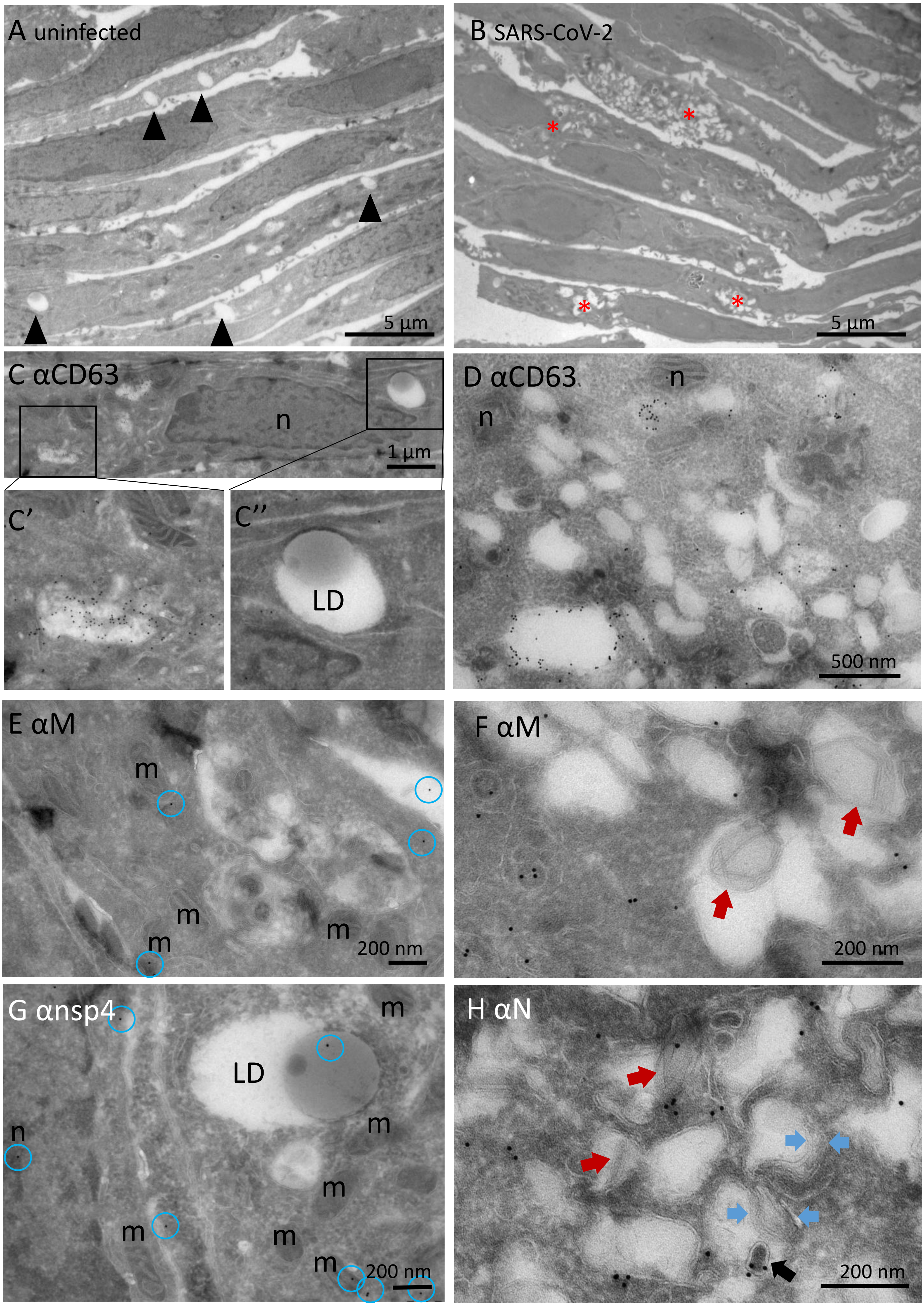

**Supplemental Figure 4. E-lucent compartment characterisation.** Electron microscopy and immuno-EM on uninfected Vero cells (left column) and in right column, SARS-CoV-2 infected Vero cells. A) Low magnification of uninfected Vero cells demonstrates multiple cells with large e-lucent compartments (black arrows). B) Low magnification of SARS-CoV-2 infected Vero cells demonstrates multiple cells with smaller and clustered e-lucent compartments (red*). C) In uninfected cells, immuno-gold labelling of CD63 on C') early endosome and C") absent on lipid-filled e-lucent compartments. D) In infected cells, immuno-gold labelling of CD63 on early endosome and not present on lipid filled e-lucent compartments. E) Control of M-protein labelling on an uninfected Vero cell showing some background label (encircled in blue). F) M-protein labelling on infected Vero cells within e-lucent compartments lipid like structures (red arrows). G) Control of nsp4 labelling on uninfected Vero cell with some background label (encircled in blue). H) Immuno-gold labelling of N-protein in e-lucent compartment, black arrow points to virus particle with N-protein. Lipid like structures represented by red arrows, double membranes by blue arrows, mitochondria by m, nuclei by n, lipid droplets by LD.

### Supplemental Figure 5

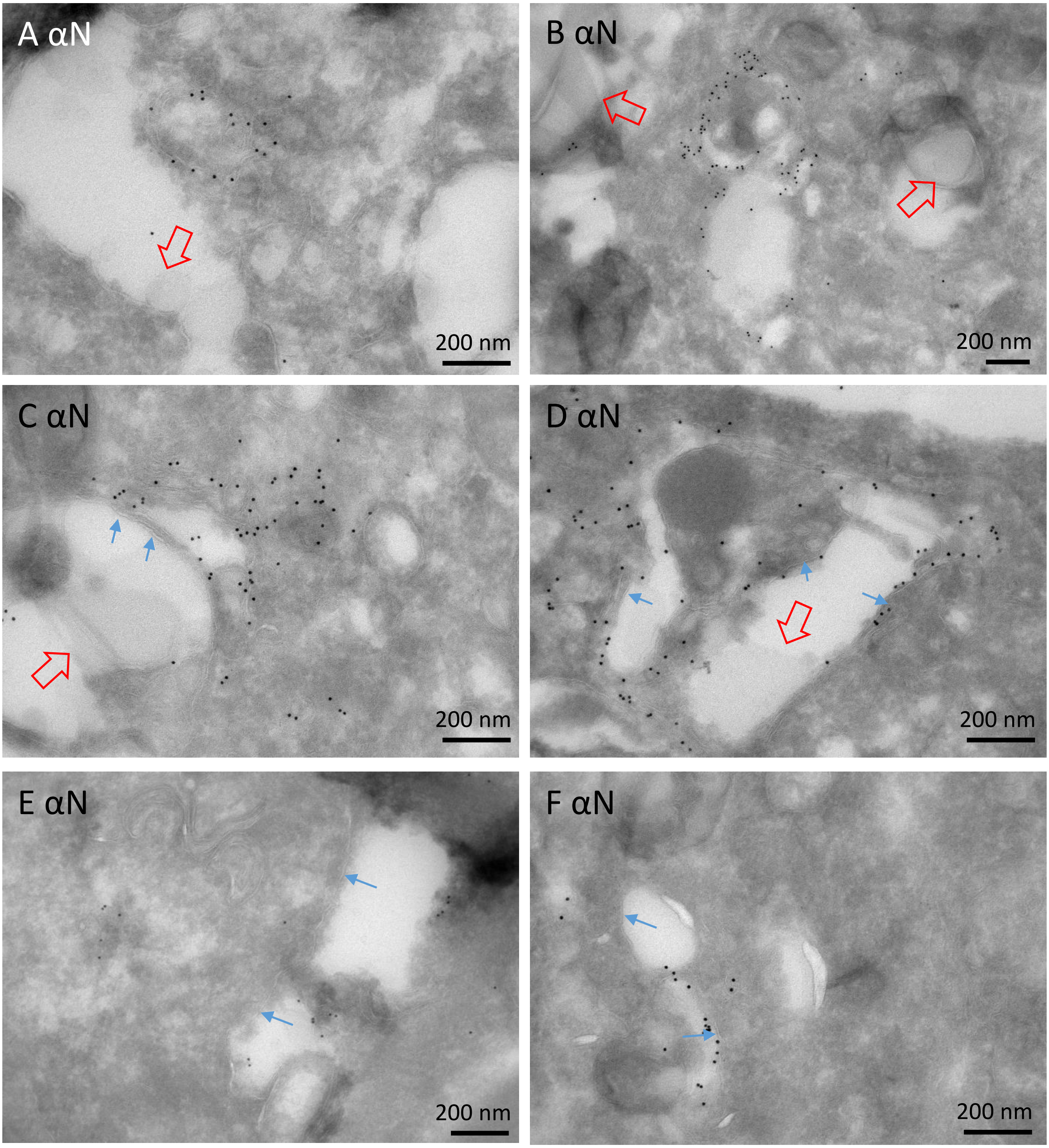

**Supplemental Figure 5. Characterization of e-lucent lipid-filled compartments in lung of COVID-19 patients.** Immuno-gold labelling with antisera raised against SARS-CoV-1 structural protein N-protein on lung tissue from COVID-19 patients (A-D patient 64 and E-F patient 58). A-D) Relatively large N-protein-positive clusters near the e-lucent compartments. E and F) Smaller N-protein-positive clusters near the e-lucent compartments. Blue arrows indicate lipid bilayer, open red arrows indicate lipid like structures.
